# The Endothelium Modulates the Prothrombotic Phenotype of Factor V Leiden: Evidence from an Ex Vivo Model

**DOI:** 10.1101/2023.11.15.567299

**Authors:** Nadine Schwarz, Jens Müller, Hannah L. McRae, Sara Reda, Behnaz Pezeshkpoor, Johannes Oldenburg, Bernd Pötzsch, Heiko Rühl

## Abstract

**Background:** Clinical expressivity of the thrombophilic factor V Leiden (FVL) mutation is highly variable. Increased activated protein C (APC) formation in response to thrombin formation has been observed in asymptomatic FVL carriers in vivo. Here we further explored this association using a recently developed endothelial colony forming cell (ECFC)-based ex-vivo model.

**Methods:** ECFCs and citrated plasma were obtained from FVL carriers with/without previous venous thromboembolism (VTE+/-, n=7 each) and seven healthy controls. Coagulation was activated by tissue factor in defibrinated recalcified plasma added to confluent cell cultures. Thrombin and APC concentration were measured over time and the respective areas under the curve (AUC) calculated. Additionally, inhibition kinetics of exogenously added APC and APC sensitivity of the prothrombinase complex were measured in plasma. Expression of thrombomodulin and endothelial protein C receptor (EPCR) on ECFCs was assessed using cell-based enzyme-linked immunosorbent assays.

**Results:** In autologous plasma on ECFCs, the APC response to thrombin formation (AUC APC/AUC thrombin), was higher in FVL VTE-than FVL VTE+ patients (0.138 versus 0.028, *P*=0.026). APC inactivation kinetics, APC sensitivity, and thrombomodulin/EPCR expression on ECFCs did not differ between these cohorts and compared to healthy controls. Cross-over experiments with plasma from FVL VTE- and FVL VTE+ patients on non-FVL ECFCs yielded indistinguishable results. In contrast, in normal plasma on FVL VTE-ECFCs the APC response remained significantly higher than on FVL VTE+ ECFCs (0.052 versus 0.022, *P*=0.011).

**Conclusions:** Consistent with results from previous in vivo experiments, APC response rates to thrombin formation were higher in asymptomatic FVL carriers compared to those with previous VTE. Our observations suggest that this increased APC response is driven by the endothelium. Further studies are warranted to elucidate yet unknown endothelial mechanisms that might modulate the clinical expressivity of FVL.

## INTRODUCTION

The factor V Leiden (FVL) mutation is the most common genetic risk factor for venous thromboembolism (VTE) with a prevalence from 3% to 15%, depending on the geographical location.^1,2^ Known under the concept of activated protein C (APC) resistance, the mutant FVL gene product lacks the APC cleavage site within its heavy chain resulting in impaired proteolytic inactivation of activated factor V (FVa)^3,4^ and, eventually in an approximately 7-fold (respectively 80-fold) increased thrombotic risk of heterozygous (respectively homozygous) carriers.^5^ However, the majority of FVL carriers remain asymptomatic throughout their lives. This highly variable clinical expressivity of FVL challenges our understanding of the interplay between genetic predisposition and molecular interactions contributing to the development of VTE.

The protein C (PC) pathway, which becomes impaired by FVL, is unique amongst anticoagulant mechanisms, since APC formation is directly linked to the amount of thrombin formed and thereby counterbalances coagulation activation with an anticoagulant response. More detailed, PC binds to the endothelial protein C receptor (EPCR) and is activated by a complex formed between thrombin and the endothelial receptor thrombomodulin (TM).^6,7^ APC, upon its release into the circulation, acts as an anticoagulant by proteolytic inactivation of FVa and activated factor VIII (FVIIIa).^8,9^ As a cofactor, circulating protein S (PS) enhances the proteolytic activity of APC.^10,11^ FVL carriers have been shown to exhibit higher thrombin and APC formation rates in vivo than non-FVL carriers after coagulation activation by low-dose recombinant activated factor VII.^12,13^ The most interesting finding in these studies was the observation that the anticoagulant APC response to thrombin formation was significantly increased in asymptomatic FVL carriers in comparison to those with a history of VTE.^13,14^

Potential explanations of this observation include alterations of plasmatic and/or endothelial components of the PC system that might modulate thrombin and APC formation rates. Among the plasmatic variables are, besides levels of involved pro- and anticoagulant proteins, other influencing factors of APC resistance^15^ and APC inactivation kinetics, the latter since elevated APC inhibitor levels have been associated with an increased thrombotic risk.^16–18^ Endothelial variables include, in particular, EPCR and TM, since their downregulation or variants in their encoding genes (*PROCR* respectively *THBD*) have been suggested as thrombotic risk factors.^19–23^

We have recently introduced an ex vivo model of the PC system, in which endothelial colony forming cells (ECFCs) and autologous plasma are utilized for personalized assessment of the functionality of the PC pathway.^19^ In the present study this experimental approach amongst others was used to further investigate plasmatic and endothelial factors that potentially modulate the APC response to thrombin in FVL carriers with or without previous VTE, in order to better understand the variable clinical expressivity of FVL.

## METHODS

The data that support the findings of this study are available from the corresponding author upon reasonable request. A list of the materials and commercially available assays used is provided in the **Supplemental Methods**. This prospective study was conducted from February 2020 through June 2023 at the Institute of Experimental Hematology and Transfusion Medicine, University Hospital Bonn, Germany. The study proposal was approved by the Institutional Review Board and Ethics Committee of the Medical Faculty of the University of Bonn. Written informed consent was obtained from all participants in compliance with the declaration of Helsinki. The procedures followed were in accordance with institutional guidelines.

### Study Participants

FVL carriers with and without history of thrombosis and FVL non-carriers were recruited from the thrombophilia outpatient clinic of our institution and from our blood donation service, respectively. Other thrombophilic risk factors including *F2* 20210G>A, deficiencies of antithrombin (AT), PC, or PS, or the presence of antiphospholipid antibodies were ruled out prior to inclusion. Molecular genetic testing for variants in *F5* (factor V, FV), *F2* (prothrombin, FII), *PROC* (PC), *PROS* (PS), *PROCR* (EPCR), *THBD* (TM) and other genes involved in the PC system was performed as described in the **Supplemental Methods**. Exclusion criteria were antiplatelet or anticoagulant medication within two weeks, and/or VTE within six months prior to blood sampling, arterial cardiovascular or malignant diseases, renal or hepatic disorders, and for female participants, pregnancy, and breast feeding. The collection and processing of blood samples is described in the **Supplemental Methods**.

### Isolation and Cultivation of Endothelial Colony Forming Cells

ECFCs were isolated and cultured as described elsewhere.^20,21^ In brief, heparinized blood was diluted with Dulbecco’s phosphate-buffered saline (DPBS) and layered over Ficoll-Paque Plus solution. Subsequently, tubes were centrifuged for 20 minutes at 1,000 x g at room temperature without brake. The buffy coat containing mononuclear cells was then collected, diluted in DPBS and centrifuged for 7 minutes at 540 x g at room temperature. The cellular pellet was resuspended in EBM^TM^-2 Endothelial Cell Growth Basal Medium-2 supplemented with the EGM^TM^-2 SingleQuots^TM^ Supplements (hydrocortisone, human basic fibroblast growth factor, vascular endothelial growth factor, R3 insulin like growth factor-1, ascorbic acid, human epidermal growth factor, gentamycin/amphotericin-1000, heparin) and 18% fetal bovine serum (FBS) and centrifuged again for 7 minutes at 540 x g at room temperature. Afterwards, cells were resuspended in the same medium and cells seeded in collagen-coated 48-well plates. After 24 hours of incubation at 37°C in a 95% air/5% CO_2_ atmosphere saturated with H_2_O, non-adherent cells and debris were removed and the medium exchanged twice weekly. The appearance of cell colonies with a cobblestone-like appearance was monitored daily after two weeks of culture. ECFC cultures were expanded by dissociation using 0.5% trypsin-ethylenediaminetetraacetic acid (EDTA) and transference to 24-well-plates, subsequently to 6-well plates, and eventually to T75 flasks, all collagen-coated. When nearing confluence, cells of six T75 flasks derived from one donor were resuspended in FBS containing 5% dimethyl sulfoxide and cryopreserved in liquid nitrogen. For experiments, frozen ECFCs were thawed, washed, and cultivated in the medium described above. All cellular assays were performed with ECFCs of passage number 7 seeded on collagen-coated 24-well, 48-well, or 96-well plates and grown until reaching confluency.

### Assessment of Endothelial Cell-Dependent Thrombin and Activated Protein C Formation

The evaluation of thrombin and APC formation in plasma or in a purified system on ECFCs was performed as described previously^19^, with modifications. For the variant using plasma, cells were washed twice with DPBS in 24-well plates and 400 µL citrated platelet-poor plasma, previously defibrinated using Batroxobin, was added to the cell culture. Thrombin and subsequent APC formation were initiated by addition of CaCl_2_, tissue factor, and phospholipids at final concentrations of 16.6 mmol/L, 1 pmol/L, and 4 µmol/L, respectively. For monitoring of thrombin and APC generation over time, reactions were stopped and active enzymes in the reaction mixture stabilized by diluting aliquots of the supernatant plasma 1:100 into phosphate-buffered saline (PBS) containing 200 µmol/L argatroban, 3 mmol/L MgCl_2_, and 0.1 % bovine serum albumin (BSA, for thrombin measurement), or 1:10 into tris-buffered saline (TBS), containing 1,000 KIU/mL aprotinin, 0.5 mg/mL bivalirudin, 1 mmol/L MgCl_2_, 7.5 mmol/L CaCl_2_, and 0.1 % BSA (APC sample buffer). All samples were stored at below -70°C until measured using oligonucleotide-enzyme-capture-assays (OECAs) which were initially described by Müller *et al*.^22,23^

### Monitoring of Activated Protein C Inactivation Kinetics in Plasma

For measurement of APC inhibition in plasma over time, human APC at a final concentration of 89 pmol/L (5 ng/mL) was added to citrated plasma and the mixture incubated at 37°C under agitation (300 rpm). Aliquots were taken over time and the reaction stopped by adding 5,000 KIU/mL aprotinin and 2,5 mg/ml bivalirudin (final concentration). Samples were stored at below -70°C until measured using the APC-OECA.

### Assessment of Activated Protein C Sensitivity Using a Prothrombinase-based Assay

To evaluate APC resistance independent from influencing factors other than factor V (FV), a prothrombinase-based assay was performed as described before^24^ with modifications. In brief, citrated plasma was diluted 1:2,000 into a phospholipid-containing buffer solution (20 mmol/L 4-(2-hydroxyethyl)-1-piperazineethanesulfonic acid, pH 7.4, 137 mmol/L NaCl, 2.5 mmol/L CaCl_2_, 5 mg/mL BSA, 5 µg/ml phospholipids) and all FV in the reaction mixture activated by incubation with 200 mU/mL Russel’s viper venom factor V activator at 37°C for 10 minutes. The mixture was divided into two parts and APC (0.64 nmol/L final concentration) was added to one part of the mixture while an equal volume of the buffer solution was added to the other part to serve as a control. Both mixtures were incubated at 37°C until reactions were stopped after 20 minutes by addition of an APC-inhibiting aptamer (HS02-52G, 10 nmol/L final concentration).^25^ Subsequently, activated factor X (FXa, 10 pmol/L), FII (10 nmol/L), and a thrombin-specific fluorogenic substrate (Boc-Asp(OBzl)-Pro-Arg-AMC, 100 µmol/L) were added and the formation of thrombin monitored over 10 minutes using a fluorescence plate reader (Synergy 2, BioTek Instruments, Bad Friedrichshall, Germany). The APC sensitivity ratio (APC-sr) was determined by dividing measured optical density (OD) to the OD of controls.

### Cell-based Enzyme Linked Immunosorbent Assay

Surface expression of EPCR and TM was measured using a cell-based enzyme-linked immunosorbent assay (cell-ELISA). ECFCs were washed and fixated with 1 % glutaraldehyde for 15 minutes on ice. Afterwards, the cells were washed and blocked with 1 % BSA for 1 hour at room temperature. A murine IgG isotype control, anti-TM antibody, or anti-EPCR antibody, labelled with horseradish peroxidase (HRP) using a HRP Conjugation Kit (Abcam, Cambridge, United Kingdom), was incubated on cells at a concentration of 0.25 µg/mL for 20 minutes at room temperature. After washing the cells twice, 3,3′,5,5′-tetramethylbenzidine substrate was added and the reaction stopped after 5 minutes. The OD at 450 nm was measured using a plate reader and background absorbance of unstained controls subtracted. Subsequently, cells were stained using Janus Green stain according to the manufacturer’s instructions and the OD of HCl-eluted stain was measured at 595 nm. To normalize for cell density, the ratio of the background-corrected OD at 450 nm and the OD at 595 nm was calculated. Calibration curves of HRP-labelled control antibody were processed in parallel covering a range from 0 to 0.5 ng/mL.

### Data Analysis

Data are generally presented as median and interquartile range (IQR). The area under the curve (AUC) was estimated using the least squares method. Normality of data was assessed using the Shapiro-Wilk test. The Kruskall-Wallis test followed by pairwise comparison using the Dunn procedure or the Mann-Whitney test were used to compare datasets. The Bonferroni method was used to correct for multiple comparisons. Two-sided, unpaired tests were used and *P* values ≤ 0.05 were considered significant. Calculations were performed using GraphPad Prism version 9.5 (GraphPad Software, Inc., San Diego, USA). N.S. had full access to all the data in the study and takes responsibility for its integrity and the data analysis.

## RESULTS

In the cohorts of asymptomatic FVL carriers (FVL VTE-), FVL carriers with a history of unprovoked VTE, (FVL VTE+), and FVL non-carriers (non-FVL), success rates of ECFC isolation and cell cultivation parameters were comparable, yielding a final study population of n=7 in each cohort, with a slightly higher number of females and of comparable age (**Table 1**).

**Table 1.** Isolation of endothelial colony forming cells and study population.

|  | Non-FVL | FVL VTE- | FVL VTE+ |
| --- | --- | --- | --- |
| Successful ECFC isolations, n (%) | 7/16 (44%) | 7/12 (58%) | 7/13 (54%) |
| ECFC colonies, n | 1.9 (1-3) | 1.4 (1-3) | 1.3 (1-3) |
| Days in culture, n | 38 (35-42) | 38 (32-48) | 39 (31-45) |
| Sex, males/females, n | 3/4 | 2/5 | 3/4 |
| Age, years, n | 44 (27-60) | 39 (22-60) | 45 (28-57) |
| FVL homozygous, n | - | 2 | 1 |
| FV HR2 haplotype, n | - | 1 | 2 |

Number of endothelial colony forming cell (ECFC) colonies, days in culture, and age are presented as mean and range. FV, factor V; FVL, factor V Leiden; VTE, venous thromboembolism.

### The Activated Protein C Response to Thrombin Formation is Increased in Asymptomatic Factor V Leiden Carriers

Thrombin and APC formation rates were assessed in the ECFC-based model of the PC pathway using autologous plasma (**Figure 1A**). In all cohorts, thrombin formation peaked at 10 minutes and levels almost fully declined after 60 minutes (**Figure 1B**) while APC peaked at 40 minutes and increased levels persisted throughout the observation time of 120 minutes (**Figure 1C**). As a measure of APC formation in response to thrombin formation, the respective AUCs were calculated for each individual subject and the ratio between the corresponding AUCs (AUC APC/AUC thrombin) was compared between cohorts. While the ratio did not differ between the non-FVL and FVL VTE+ subgroups (Kruskall-Wallis test *P*>0.999), it was found to be 3-fold higher in asymptomatic FVL VTE-compared to non-FVL carriers (median, IQR of 0.138, 0.110-0.151 versus 0.047, 0.025-0.054; *P*=0.007) and 5-fold higher compared to the FVL VTE+ group (0.028, 0.022-0.092; *P*=0.026) (**Figure 1D**). Of note, homozygous FVL carriers showed higher AUCs of both thrombin and APC formation than heterozygous FVL carriers within the FVL VTE-group (AUC thrombin: >2209 versus <1775 nmol·min·L^-1^, AUC APC: >305 versus <265 nmol·min·L^-1^) and the FVL VTE+ group (AUC thrombin: 3441 versus <3269 nmol·min·L^-1^, AUC APC: 286 versus <208 nmol·min·L^-1^), but the ratio between AUCs lay near the median of the respective groups (**Table S1, Figure 1F**).

**Figure 1.**
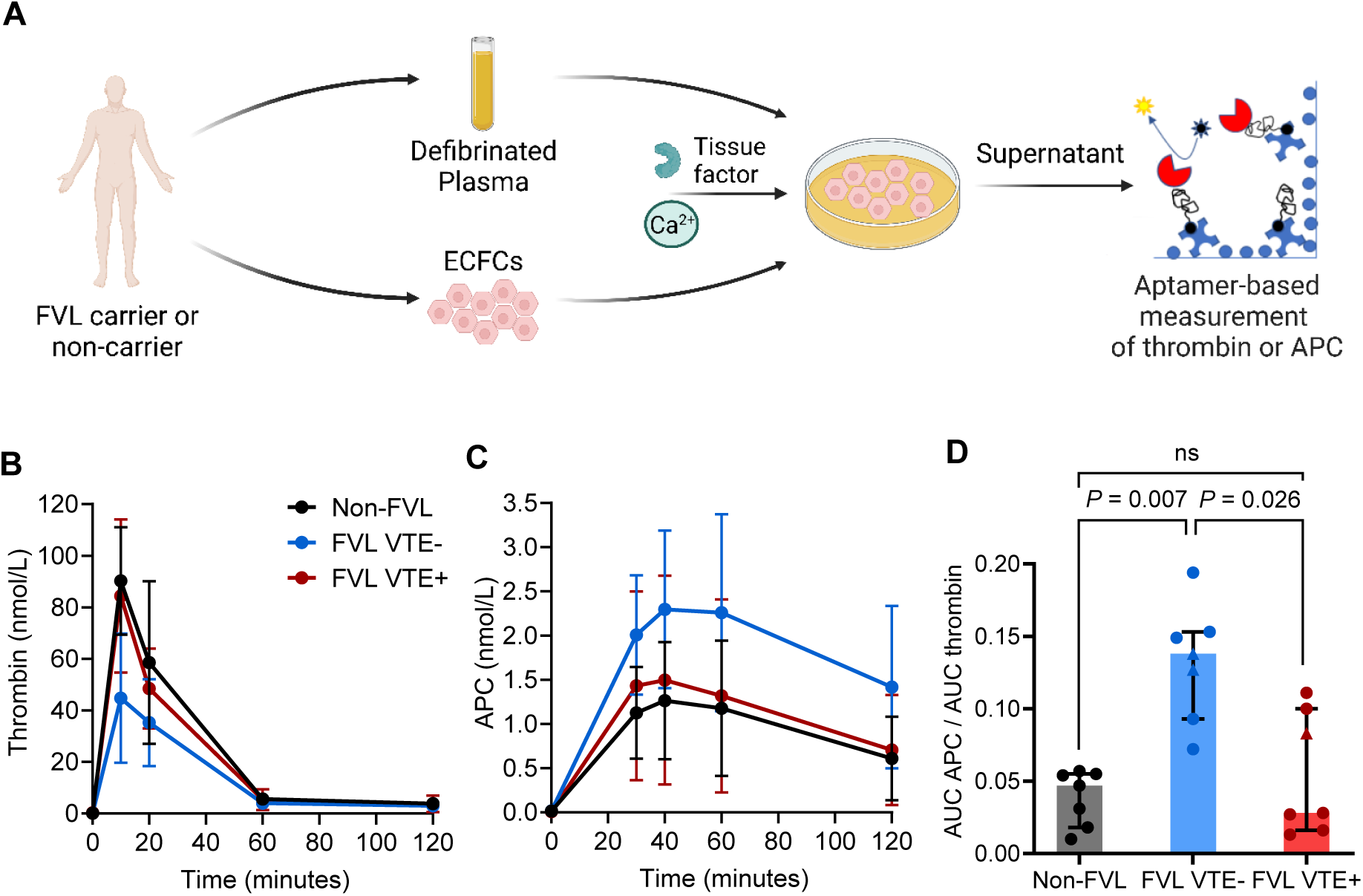
Protein C pathway modelling using endothelial cells and autologous plasma in factor V Leiden carriers and healthy controls. **A**, To assess the endothelial activated protein C (APC) formation capacity, confluent monolayers of endothelial colony forming cells (ECFCs) were overlaid with autologous defibrinated, citrated plasma and thrombin formation induced by addition of tissue factor and CaCl_2_. In the supernatant, time-dependent formation of (**B**) thrombin and (**C**) APC were monitored in non-factor V Leiden (FVL) carriers (depicted in black), FVL carriers with a history of venous thromboembolism (VTE, FVL VTE+, red) or asymptomatic FVL carriers (FVL VTE-, blue, n=7 each). **D**, The ratio between the area under the curve (AUC) of thrombin formation and the AUC of APC formation (AUC APC/AUC thrombin) as measure of the APC response were compared between cohorts using the Kruskall-Wallis test followed by pairwise comparison using the Dunn procedure. Data are shown as median and interquartile range. Measurement results in homozygous FVL carriers are distinguished by triangles. ns, not significant.

In addition to FVL, other gene variants were detected in the study population, including *F5* 6755A>G (indicative for the FV HR2 haplotype), *THBD* 1418C>T, *PROCR* 4600A>G, and *PROCR* 4678G>C. Their allele frequencies in the FVL VTE-, FVL VTE+, and non-FVL carriers are listed in **Table S1**, along with the respective ranges of the APC response (AUC APC/AUC thrombin). These ranges in homozygous *PROCR* 4678G>C carriers, and in any carriers of other gene variants lay within the ranges observed in respective non-carriers in both FVL cohorts (**Table S1**).

### Examinations of Plasmatic Determinants Potentially Influencing the Activated Protein C Response

To study whether plasmatic variables might be causative for an increased APC response in asymptomatic FVL carriers, we compared coagulation factor and inhibitor levels, the plasmatic APC-inhibitory capacity, and APC resistance between the FVL VTE-, FVL VTE+, and non-FVL cohort. Plasma levels of coagulation factors (FII, FV, factor VII, factor VIII, factor X, and factor XI) and inhibitors (AT, PC, and PS) did not differ statistically significantly between cohorts (Kruskall-Wallis test *P*>0.05) (**Figure 2A**). The APC half-life was found to be virtually identical (Kruskall-Wallis test *P*>0.05) when APC was added to plasma and decline of the enzyme was followed over time, with median (IQR) of 19.2 (18.8-20.7) minutes in non-FVL, 19.8 (17.4-21.5) minutes in FVL VTE-, and 20.9 (19.4-21.8) minutes in FVL VTE+ cohorts (**Figure 2B**). As a measure of APC resistance, we performed a prothrombinase-based assay in diluted plasma under targeted FV-activation and determined the APC-sr via measurement of thrombin formation over time with and without incubation with APC. As expected, the median (IQR) APC-sr in the non-FVL cohort (22.9, 17.8-22.9%) was significantly lower when compared to the FVL VTE-subgroup (49.4, 47.7-64.4%; Kruskall Wallis test *P*=0.012) and to the FVL VTE+ subgroup (54.0, 52.4-57.3%; *P*=0.002) whereas the APC-sr did not differ between FVL VTE- and FVL VTE+ (*P*>0.999) (**Figure 2C**). The APC-sr also showed no difference between FVL VTE- and FVL VTE+, when homozygous FVL carriers, who showed a higher APC-sr than heterozygous carriers, were excluded (*P*=0.636).

**Figure 2.**
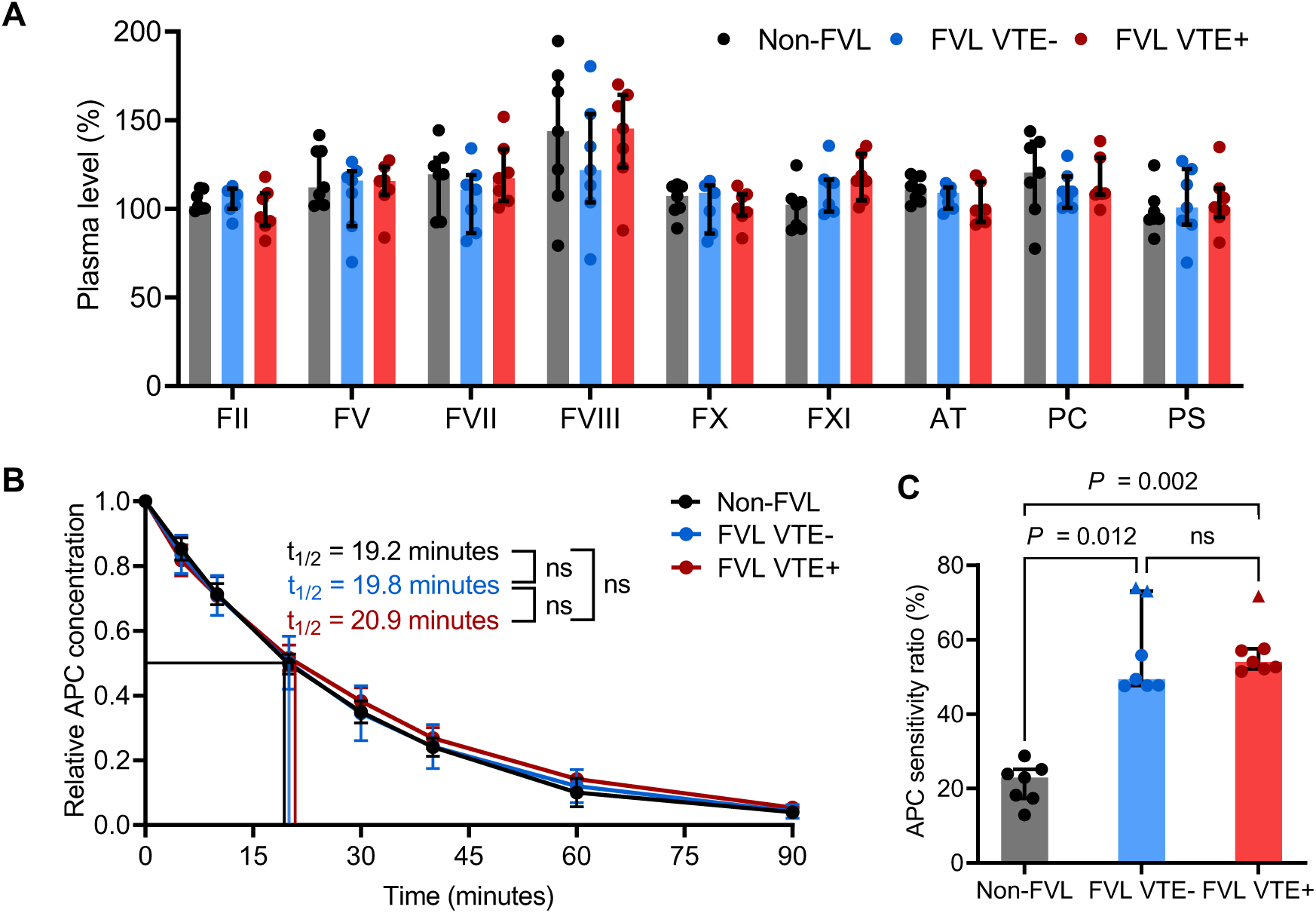
Evaluation of plasmatic factors potentially affecting the activated protein C response. Data obtained in non-factor V Leiden (FVL) carriers (depicted in black), FVL carriers with a history of venous thromboembolism (VTE, FVL VTE+, red) or asymptomatic FVL carriers (FVL VTE-, blue, n=7 each) are shown as median and interquartile range. **A**, Plasma levels of coagulation factors and inhibitors were compared (*P*>0.05 each). **B**, Activated protein C (APC) inhibition kinetics were evaluated by addition of APC (5 ng/mL final concentration) to citrated plasma and monitoring of residual plasma concentrations over time by an oligonucleotide-based enzyme capture assay. The calculated half-life time (t_1/2_) was compared between cohorts. **C**, The APC resistance of activated factor V (FVa) was assessed by measuring the residual prothrombinase activity after inactivation by APC and the APC sensitivity ratio was determined. Data are shown as median and interquartile range. Measurement results in homozygous FVL carriers are depicted by triangles. Comparisons between cohorts were performed using the Kruskall-Wallis test followed by pairwise comparison using the Dunn procedure. AT, antithrombin; FII-XI, factor II-XI; ns, not significant; PS, protein S.

Modifying the autologous PC pathway model with focus on plasmatic variability, FVL VTE- or FVL VTE+ plasma from each study participant was added to the same non-FVL ECFC culture and the APC response was evaluated via the AUC APC/AUC thrombin ratio. In contrast to the results from the autologous system, the APC response did not differ statistically significantly between the FVL VTE- and FVL VTE+ groups (Mann-Whitney test *P*=0.053), which suggests that differences in the autologous approach are not driven by plasmatic variation (**Figure 3A**).

**Figure 3.**
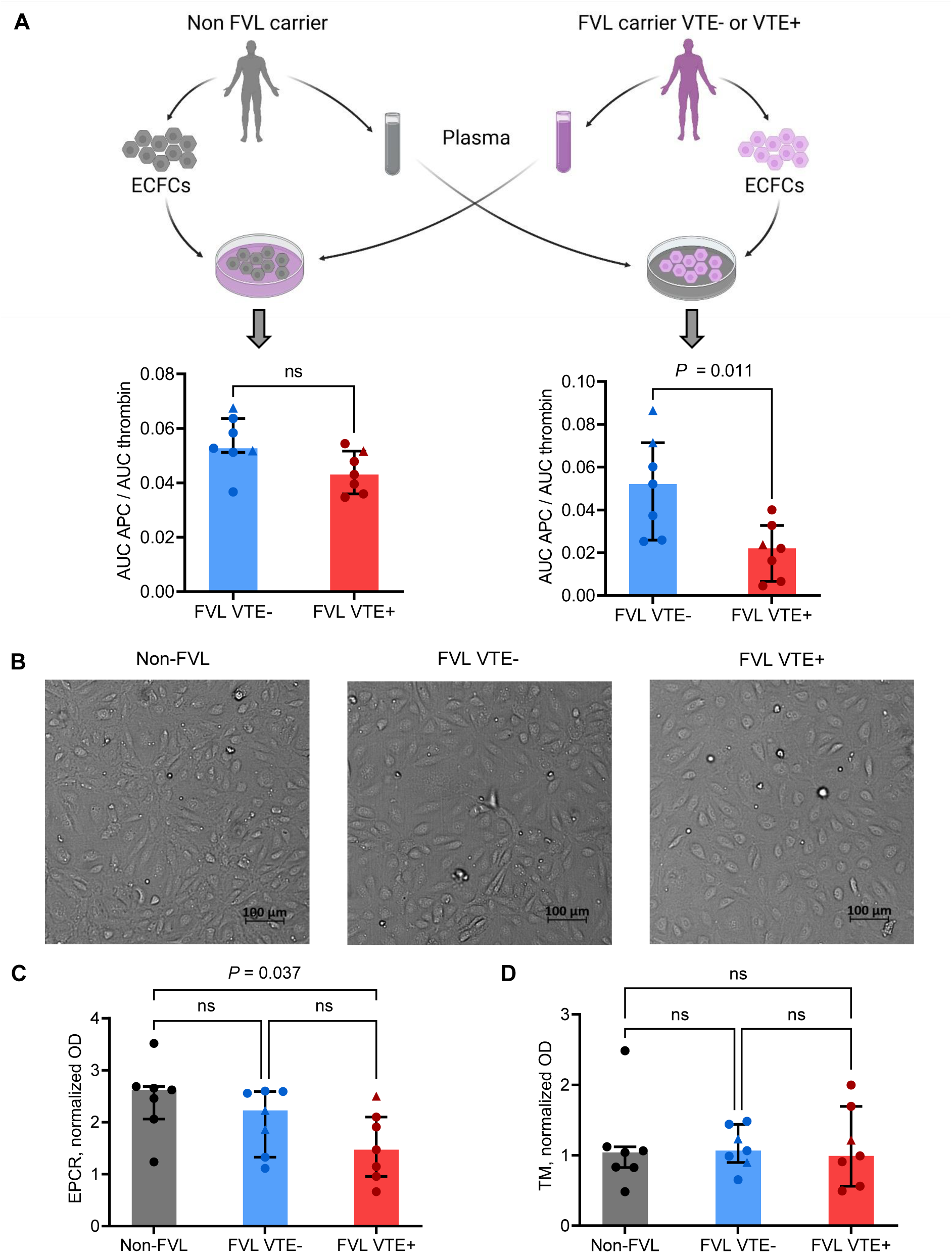
Evaluation of endothelial factors potentially affecting the activated protein C response. **A**, The activated protein C (APC) response was studied in the endothelial cell-based protein C pathway model using either endothelial colony-forming cells (ECFCs) from a non-FVL carriers and plasma from FVL VTE- and FVL VTE+ individuals; or ECFCs from FVL VTE- and FVL VTE+ individuals and pooled normal plasma. The ratio between the area under the curve (AUC) of APC formation (AUC APC) and the AUC of thrombin formation (AUC thrombin) as measure of the APC response was compared between cohorts using the Mann Whitney test. **B**, Representative micrographs of endothelial colony forming cells (ECFCs) from a non-factor V Leiden (FVL) carrier, a FVL carrier with a history of venous thromboembolism (VTE, FVL VTE+), and an asymptomatic FVL carrier (FVL VTE-) are shown (Axio Observer, camera Axiocam 702 mono (Carl Zeiss Microscopy). The following data, obtained in non-FVL (black), FVL VTE-(blue), and FVL VTE+ (red) individuals (n=7 each), are shown as median and interquartile range, and were compared using the Kruskall-Wallis test followed by pairwise comparison using the Dunn procedure. Measurement results in homozygous FVL carriers are distinguished by triangles. Expression of (**C**) endothelial protein C receptor (EPCR) and (**D**) thrombomodulin (TM) on ECFCs was studied by cell-based enzyme-linked immunosorbent assays, in which ECFCs were fixated and incubated with the respective antibody. The background corrected optical density (OD) was normalized for cell density assessed by Janus Green staining. ns, not significant.

### Endothelial-dependent Activated Protein C response in Healthy Control Plasma is Higher in Asymptomatic Factor V Leiden Carriers

Modifying the autologous PC pathway model with focus on endothelial variability, the same pooled normal plasma was added to ECFC cultures from FVL VTE- or FVL VTE+ individuals and the APC response was evaluated via the AUC APC/AUC thrombin ratio. As measured in the autologous system and in contrast to the varying plasma approach, the APC response in asymptomatic FVL VTE-carriers was found to be significantly higher compared to the FVL VTE+ subgroup in this endothelial-dependent approach (Mann-Whitney test *P*=0.011) (**Figure 3A**). This observation suggests that the increased APC response in asymptomatic FVL carriers is driven by the endothelium rather than by plasma components.

For evaluation of this phenomenon, we compared characteristics of ECFCs obtained from non-FVL, FVL VTE-, and FVL VTE+ carriers and further studied individual endothelial factors with potential influence on the APC response. ECFC morphology in light microscopy (**Figure 3B, Figure S1**), total protein amount of a confluent colony (**Figure S2A**), and characteristic surface marker expression in flow cytometry (**Figure S2B**) did not differ between cohorts. The surface expression of EPCR and TM on ECFCs was quantitatively measured by cell-ELISA. EPCR expression was found to be higher in non-FVL carriers compared to the FVL VTE+ subgroup (Kruskall-Wallis test *P*=0.037) but did not differ between non-FVL and FVL VTE-(*P*=0.635), and between FVL VTE- and FVL VTE+ (*P*=0.635) (**Figure 3C**). In addition, we found that TM expression on the cell surface did not differ between cohorts (Kruskall-Wallis test *P*>0.05) (**Figure 3C**).

## DISCUSSION

In the common understanding on how FVL affects thrombotic risk, thrombogenicity is driven by the prolonged half-life of FVa, resulting in increased prothrombinase complex activity. Accordingly, the thrombotic risk is higher in homozygous FVL carriers, who have higher levels of APC-resistant FVa, than in heterozygous carriers.^5^ Provided that the endothelial PC pathway is intact, higher thrombin formation rates should result in higher APC formation rates. Indeed, using an ECFC-based ex vivo model of the PC pathway, we have recently shown that the APC response to thrombin formation was increased in FVL carriers, the majority of whom had no history of VTE.^19^ In the present study, using the same model but distinguishing between symptomatic and asymptomatic FVL carriers, we observed an increased APC response only in the latter cohort. Of note, while homozygous FVL carriers showed the highest AUCs of thrombin in their respective cohorts (VTE+ and VTE-) and the highest AUCs of APC overall, the ratio of AUC APC/AUC thrombin in homozygous FVL carriers lay near the median of their respective cohorts. This can be explained by the ex vivo model being a dynamic system, in which APC formation downregulates thrombin formation as does the PC pathway itself. The ex vivo results are in line with previous in vivo findings of increased APC response rates after injection of rFVIIa in asymptomatic FVL carriers compared to those with a history of VTE.^13^ While this further strengthens the validity of the ECFC-based model of the PC pathway, the question remains which factors are causative for the observed differences in the APC response.

To narrow down potentially influencing factors, additional experiments were conducted using a single ECFC line from a non-FVL carrier and plasma from FVL carriers with and without a history of VTE. As a result, the differences in the APC response between FVL VTE+ and FVL VTE-disappeared, speaking against a role of plasmatic factors in the observed differences in the APC response. Further investigation of potential influencing factors in plasma revealed no differences in coagulation factors and inhibitors, APC inhibition kinetics, or APC resistance between the cohorts of FVL carriers. The APC response in carriers of the FV HR2 haplotype (one in FVL VTE-, two in FVL+), which is known to modulate APC resistance in FVL carriers,^26,27^ lay within the ranges of the APC resistance in both cohorts. To sum up, the observed differences in the APC response could not be explained by the examined plasmatic variables.

When ECFC lines from the FVL VTE+ and FVL VTE-cohorts were combined with normal pooled plasma in the ECFC model, the differences in the APC response observed in the autologous approach, could be reproduced, albeit at a lesser extent. Taken together, the obtained data using the ECFC-based model of the PC pathway suggest that endothelial factors modulate the APC response.

Well established endothelial determinants of APC formation are the receptors TM and EPCR. Since PC activation is directly linked to the amount of TM and EPCR presented on the endothelial cell surface,^28,29^ we studied their expression on ECFCs using cell-ELISAs. Although we observed no difference in TM or EPCR expression between asymptomatic and symptomatic FVL carriers in a resting state, the localization of receptors across the endothelial cell membrane could change upon coagulation activation. While they are usually presented on the cell surface, TM and EPCR can be endocytosed in response to stimulation making them unavailable for further ligand binding and, at the same time, bound ligands are cleared from the circulation. Thrombin induces the internalization of TM which is followed by degradation of thrombin.^30–32^ Similarly, ligand binding to EPCR promotes endocytosis of the receptor. In vitro studies showed that PC, APC, FVII, and FVIIa bind to EPCR with a similar affinity^33^ and are internalized via endocytosis in a similar rate.^34^ Therefore, EPCR endocytosis downregulates both the anticoagulant response of the PC pathway and the procoagulant response of the extrinsic pathway. While in theory, internalization of endothelial receptors should have a proportionate effect on pro- and anticoagulant mechanisms, one might speculate if a dysbalanced internalization could explain the observed differences in the APC response.

Gene variants in *THBD* or *PROCR* have also been postulated to affect APC formation and thrombotic risk. Navarro et al^35^ reported an association of the *THBD* c.1418 C>T polymorphism with reduced VTE risk and increased circulating APC levels in carriers of the 1418T allele. In addition, they performed functional studies in cultured human umbilical vein endothelial cells and observed increased PC activation on cells carrying the 1418T allele. Furthermore, different *PROCR* haplotypes and their functional phenotypes have been characterized. The H1 (or A1) haplotype, tagged by the 4678 G>C sequence, was associated with reduced EPCR shedding and decreased VTE risk.^36,37^ Medina et al^38^ showed that FVL carriers with the *PROCR* H1 haplotype have a reduced thrombotic risk, although no association between APC levels and the polymorphism could be established. In opposite, the H3 (or A3) haplotype, tagged by the 4600 A>G sequence, was associated with lower PC activation, increased EPCR shedding and higher VTE risk.^39,40^ While the association between an increased thrombotic risk and the *PROCR* 4600 A>G variant was confirmed by larger trials and meta-analyses,^41–43^ these studies did not find an association between the common *THBD* c.1418C>T polymorphism^44–47^ and the *PROCR* 4678 G>C variant^39,41^ and reduced risk of VTE. In our study, the APC response in symptomatic and asymptomatic FVL carriers, who were carriers of the aforementioned *THBD* or *PROCR* variants, lay within the range of the APC response in the respective cohorts. Therefore, they cannot explain the observed differences in the APC response. However, a potential effect *THBD* or *PROCR* variants on the APC response cannot be excluded but would require examination in study populations who would need to be accordingly selected.

In conclusion, our findings confirm previous in vivo evidence that the extent of APC formation in response to thrombin modulates the thrombotic risk in FVL and might therefore at least partially explain the variable clinical expressivity of this thrombophilic mutation. Furthermore, they underline the validity of the utilized ECFC-based ex vivo model in assessing the functionality of the PC pathway regarding clinically relevant endpoints on a personalized level. Although the underlying mechanism remains yet to be identified, the obtained data strongly suggest that endothelial variables are a major driver of variation in the APC response in symptomatic and asymptomatic FVL carriers. In addition to further elucidation of causative factors, further studies are warranted to examine a potential role of the endothelium in other types of thrombophilia.

## Non-standard Abbreviations and Acronyms

APC: activated protein C;
APC-sr: APC sensitivity ratio;
AUC: area under the curve;
AT: antithrombin;
BSA: bovine serum albumin;
cell-ELISA: cell-based enzyme-linked immunosorbent assay;
DPBS: Dulbecco’s phosphate-buffered saline;
ECFC: endothelial colony forming cell;
EDTA: ethylenediaminetetraacetic acid;
EPCR: endothelial protein C receptor;
FBS: fetal bovine serum;
FII: prothrombin;
FIX: factor IX;
FV: factor V;
FVa: activated factor V;
FVII: factor VII;
FVIII: factor VIII;
FVIIIa: activated factor VIII;
FVL: factor V Leiden;
FX: factor X;
FXa: activated factor X;
FXI: factor XI;horseradish peroxidase (HRP);
IQR: interquartile range;
OECA: oligonucleotide-based enzyme capture assay;
OD: optical density;
PBS: phosphate-buffered saline;
PC: protein C;
PS: protein S;
TBS: Tris buffered saline;
TM: thrombomodulin;
VTE: venous thromboembolism.

## Acknowledgements

Figure 1A and 3A have been created with BioRender.com.

## Supplemental Material

Supplemental Methods

Table S1

Figure S1-S2

Major Resources Table

## NOVELTY AND SIGNIFICANCE

### What is known?

- The factor V Leiden (FVL) mutation impairs inactivation of procoagulant factor V by anticoagulant activated protein C (APC).
- FVL is the most prevalent hereditary risk factor for venous thromboembolism (VTE) with a highly variable clinical expressivity.
- In vivo, asymptomatic FVL carriers have been shown to form more APC upon coagulation activation than those with previous VTE.

### What new information does this article contribute?

- An increased APC response in asymptomatic FVL carriers was confirmed in an ex vivo model using patient-specific endothelial cells and autologous plasma.
- Differences in the APC response were driven by the patients’ endothelial cells rather than variables in plasma.
- Endothelial mechanisms might modulate the thrombotic risk in FVL carriers.

The FVL mutation alters the site at which this procoagulant plasma protein is cleaved by the anticoagulant APC, which is formed on the endothelial surface upon coagulation activation. While it is the most common hereditary thrombophilia, its clinical phenotype – venous thrombosis and pulmonary embolism – shows a highly variable expressivity. Recently, we have shown in vivo, that asymptomatic FVL carriers form more APC after extrinsic coagulation activation than those with a history of VTE. Using an ex vivo model based on patient-specific endothelial cells and autologous plasma we were able to confirm these differences in the APC response, that depended on the clinical phenotype, in FVL carriers. Selective analysis of endothelial and plasmatic variables that might affect APC formation we found that the increased APC formation in asymptomatic FVL carriers was driven by the patients’ endothelium. These data suggest that endothelial mechanisms might modulate the thrombotic risk in FVL carriers.

## REFERENCES

1. Rees DC, Cox M, Clegg JB. World distribution of factor V Leiden. Lancet. 1995;346:1133–1134. doi: 10.1016/s0140-6736(95)91803-5.

2. Rosendaal FR, Doggen CJ, Zivelin A, Arruda VR, Aiach M, Siscovick DS, Hillarp A, Watzke HH, Bernardi F, Cumming AM, et al. Geographic distribution of the 20210 G to A prothrombin variant. Thromb Haemost. 1998;79:706–708.

3. Dahlbäck B, Hildebrand B. Inherited resistance to activated protein C is corrected by anticoagulant cofactor activity found to be a property of factor V. Proc Natl Acad Sci U S A. 1994;91:1396–1400. doi: 10.1073/pnas.91.4.1396.

4. Bertina RM, Koeleman BP, Koster T, Rosendaal FR, Dirven RJ, Ronde H de, van der Velden PA, Reitsma PH. Mutation in blood coagulation factor V associated with resistance to activated protein C. Nature. 1994;369:64–67. doi: 10.1038/369064a0.

5. Rosendaal FR, Koster T, Vandenbroucke JP, Reitsma PH. High risk of thrombosis in patients homozygous for factor V Leiden (activated protein C resistance). Blood. 1995;85:1504–1508.

6. Esmon CT. The roles of protein C and thrombomodulin in the regulation of blood coagulation. J Biol Chem. 1989;264:4743–4746.

7. Taylor FB, Peer GT, Lockhart MS, Ferrell G, Esmon CT. Endothelial cell protein C receptor plays an important role in protein C activation in vivo. Blood. 2001;97:1685–1688. doi: 10.1182/blood.v97.6.1685.

8. Walker FJ, Chavin SI, Fay PJ. Inactivation of factor VIII by activated protein C and protein S. Arch Biochem Biophys. 1987;252:322–328. doi: 10.1016/0003-9861(87)90037-3.

9. Kalafatis M, Rand MD, Mann KG. The mechanism of inactivation of human factor V and human factor Va by activated protein C. J Biol Chem. 1994;269:31869–31880.

10. Shen L, Dahlbäck B. Factor V and protein S as synergistic cofactors to activated protein C in degradation of factor VIIIa. J Biol Chem. 1994;269:18735–18738.

11. Dahlbäck B. Factor V and protein S as cofactors to activated protein C. Haematologica. 1997;82:91–95.

12. Rühl H, Winterhagen FI, Berens C, Müller J, Oldenburg J, Pötzsch B. In vivo thrombin generation and subsequent APC formation are increased in factor V Leiden carriers. Blood. 2018;131:1489– 1492. doi: 10.1182/blood-2017-12-823831.

13. Rühl H, Berens C, Winterhagen FI, Reda S, Müller J, Oldenburg J, Pötzsch B. Increased Activated Protein C Response Rates Reduce the Thrombotic Risk of Factor V Leiden Carriers But Not of Prothrombin 20210GA Carriers. Circ Res. 2019;125:523–534. doi: 10.1161/CIRCRESAHA.119.315037.

14. Rühl H, Friemann AM, Reda S, Schwarz N, Winterhagen FI, Berens C, Müller J, Oldenburg J, Pötzsch B. Activated Factor XI is Increased in Plasma in Response to Surgical Trauma but not to Recombinant Activated FVII-Induced Thrombin Formation. J Atheroscler Thromb. 2022;29:82–98. doi: 10.5551/jat.59873.

15. Castoldi E, Rosing J. APC resistance: biological basis and acquired influences. J Thromb Haemost. 2010;8:445–453. doi: 10.1111/j.1538-7836.2009.03711.x.

16. Meijers JCM, Marquart JA, Bertina RM, Bouma BN, Rosendaal FR. Protein C inhibitor (plasminogen activator inhibitor-3) and the risk of venous thrombosis. Br J Haematol. 2002;118:604–609. doi: 10.1046/j.1365-2141.2002.03652.x.

17. Basil N, Ekström M, Piitulainen E, Lindberg A, Rönmark E, Jehpsson L, Tanash H. Severe alpha-1-antitrypsin deficiency increases the risk of venous thromboembolism. J Thromb Haemost. 2021;19:1519–1525. doi: 10.1111/jth.15302.

18. Beheiri A, Langer C, Düring C, Krümpel A, Thedieck S, Nowak-Göttl U. Role of elevated alpha2-macroglobulin revisited: results of a case-control study in children with symptomatic thromboembolism. J Thromb Haemost. 2007;5:1179–1184. doi: 10.1111/j.1538-7836.2007.02534.x.

19. Schwarz N, Müller J, Yadegari H, McRae HL, Reda S, Hamedani NS, Oldenburg J, Pötzsch B, Rühl H. Ex Vivo Modeling of the PC (Protein C) Pathway Using Endothelial Cells and Plasma: A Personalized Approach. Arterioscler Thromb Vasc Biol. 2023;43:109–119. doi: 10.1161/ATVBAHA.122.318433.

20. Martin-Ramirez J, Hofman M, van den Biggelaar M, Hebbel RP, Voorberg J. Establishment of outgrowth endothelial cells from peripheral blood. Nat Protoc. 2012;7:1709–1715. doi: 10.1038/nprot.2012.093.

21. Ormiston ML, Toshner MR, Kiskin FN, Huang CJZ, Groves E, Morrell NW, Rana AA. Generation and Culture of Blood Outgrowth Endothelial Cells from Human Peripheral Blood. J Vis Exp. 2015:e53384. doi: 10.3791/53384.

22. Müller J, Becher T, Braunstein J, Berdel P, Gravius S, Rohrbach F, Oldenburg J, Mayer G, Pötzsch B. Profiling of active thrombin in human blood by supramolecular complexes. Angew Chem Int Ed Engl. 2011;50:6075–6078. doi: 10.1002/anie.201007032.

23. Müller J, Friedrich M, Becher T, Braunstein J, Kupper T, Berdel P, Gravius S, Rohrbach F, Oldenburg J, Mayer G, et al. Monitoring of plasma levels of activated protein C using a clinically applicable oligonucleotide-based enzyme capture assay. J Thromb Haemost. 2012;10:390–398. doi: 10.1111/j.1538-7836.2012.04623.x.

24. Nicolaes GA, Thomassen MC, van Oerle R, Hamulyak K, Hemker HC, Tans G, Rosing J. A prothrombinase-based assay for detection of resistance to activated protein C. Thromb Haemost. 1996;76:404–410.

25. Müller J, Isermann B, Dücker C, Salehi M, Meyer M, Friedrich M, Madhusudhan T, Oldenburg J, Mayer G, Pötzsch B. An exosite-specific ssDNA aptamer inhibits the anticoagulant functions of activated protein C and enhances inhibition by protein C inhibitor. Chem Biol. 2009;16:442–451. doi: 10.1016/j.chembiol.2009.03.007.

26. Visser MC de, Guasch JF, Kamphuisen PW, Vos HL, Rosendaal FR, Bertina RM. The HR2 haplotype of factor V: effects on factor V levels, normalized activated protein C sensitivity ratios and the risk of venous thrombosis. Thromb Haemost. 2000;83:577–582.

27. Bossone A, Cappucci F, D’Andrea G, Brancaccio V, Cibelli G, Iannaccone L, Grandone E, Margaglione M. The factor V (FV) gene ASP79HIS polymorphism modulates FV plasma levels and affects the activated protein C resistance phenotype in presence of the FV Leiden mutation. Haematologica. 2003;88:286–289.

28. Stearns-Kurosawa DJ, Kurosawa S, Mollica JS, Ferrell GL, Esmon CT. The endothelial cell protein C receptor augments protein C activation by the thrombin-thrombomodulin complex. Proc Natl Acad Sci U S A. 1996;93:10212–10216. doi: 10.1073/pnas.93.19.10212.

29. Isermann B, Hendrickson SB, Zogg M, Wing M, Cummiskey M, Kisanuki YY, Yanagisawa M, Weiler H. Endothelium-specific loss of murine thrombomodulin disrupts the protein C anticoagulant pathway and causes juvenile-onset thrombosis. J Clin Invest. 2001;108:537–546. doi: 10.1172/JCI13077.

30. Maruyama I, Majerus PW. The turnover of thrombin-thrombomodulin complex in cultured human umbilical vein endothelial cells and A549 lung cancer cells. Endocytosis and degradation of thrombin. J Biol Chem. 1985;260:15432–15438.

31. Conway EM, Boffa MC, Nowakowski B, Steiner-Mosonyi M. An ultrastructural study of thrombomodulin endocytosis: internalization occurs via clathrin-coated and non-coated pits. J Cell Physiol. 1992;151:604–612. doi: 10.1002/jcp.1041510321.

32. Teasdale MS, Bird CH, Bird P. Internalization of the anticoagulant thrombomodulin is constitutive and does not require a signal in the cytoplasmic domain. Immunol Cell Biol. 1994;72:480–488. doi: 10.1038/icb.1994.72.

33. Ghosh S, Pendurthi UR, Steinoe A, Esmon CT, Rao LVM. Endothelial cell protein C receptor acts as a cellular receptor for factor VIIa on endothelium. J Biol Chem. 2007;282:11849–11857. doi: 10.1074/jbc.M609283200.

34. Nayak RC, Sen P, Ghosh S, Gopalakrishnan R, Esmon CT, Pendurthi UR, Rao LVM. Endothelial cell protein C receptor cellular localization and trafficking: potential functional implications. Blood. 2009;114:1974–1986. doi: 10.1182/blood-2009-03-208900.

35. Navarro S, Medina P, Bonet E, Corral J, Martínez-Sales V, Martos L, Rivera M, Roselló-Lletí E, Alberca I, Roldán V, et al. Association of the thrombomodulin gene c.1418CT polymorphism with thrombomodulin levels and with venous thrombosis risk. Arterioscler Thromb Vasc Biol. 2013;33:1435–1440. doi: 10.1161/ATVBAHA.113.301360.

36. Medina P, Navarro S, Estellés A, Vayá A, Woodhams B, Mira Y, Villa P, Migaud-Fressart M, Ferrando F, Aznar J, et al. Contribution of polymorphisms in the endothelial protein C receptor gene to soluble endothelial protein C receptor and circulating activated protein C levels, and thrombotic risk. Thromb Haemost. 2004;91:905–911. doi: 10.1160/TH03-10-0657.

37. Medina P, Navarro S, Bonet E, Martos L, Estellés A, Bertina RM, Vos HL, España F. Functional analysis of two haplotypes of the human endothelial protein C receptor gene. Arterioscler Thromb Vasc Biol. 2014;34:684–690. doi: 10.1161/ATVBAHA.113.302518.

38. Medina P, Navarro S, Estellés A, Vayá A, Bertina RM, España F. Influence of the 4600A/G and 4678G/C polymorphisms in the endothelial protein C receptor (EPCR) gene on the risk of venous thromboembolism in carriers of factor V Leiden. Thromb Haemost. 2005;94:389–394. doi: 10.1160/TH05-02-0089.

39. Uitte de Willige S, van Marion V, Rosendaal FR, Vos HL, Visser MCH de, Bertina RM. Haplotypes of the EPCR gene, plasma sEPCR levels and the risk of deep venous thrombosis. J Thromb Haemost. 2004;2:1305–1310. doi: 10.1046/j.1538-7836.2004.00855.x.

40. Saposnik B, Reny J-L, Gaussem P, Emmerich J, Aiach M, Gandrille S. A haplotype of the EPCR gene is associated with increased plasma levels of sEPCR and is a candidate risk factor for thrombosis. Blood. 2004;103:1311–1318. doi: 10.1182/blood-2003-07-2520.

41. Manderstedt E, Halldén C, Lind-Halldén C, Elf J, Svensson PJ, Engström G, Melander O, Baras A, Lotta LA, Zöller B. Thrombotic Risk Determined by Protein C Receptor (PROCR) Variants among Middle-Aged and Older Adults: A Population-Based Cohort Study. Thromb Haemost. 2022;122:1326–1332. doi: 10.1055/a-1738-1564.

42. Dennis J, Johnson CY, Adediran AS, Andrade M de, Heit JA, Morange P-E, Trégouët D-A, Gagnon F. The endothelial protein C receptor (PROCR) Ser219Gly variant and risk of common thrombotic disorders: a HuGE review and meta-analysis of evidence from observational studies. Blood. 2012;119:2392–2400. doi: 10.1182/blood-2011-10-383448.

43. Stacey D, Chen L, Stanczyk PJ, Howson JMM, Mason AM, Burgess S, MacDonald S, Langdown J, McKinney H, Downes K, et al. Elucidating mechanisms of genetic cross-disease associations at the PROCR vascular disease locus. Nat Commun. 2022;13:1222. doi: 10.1038/s41467-022-28729-3.

44. Ahmad A, Sundquist K, Zöller B, Svensson PJ, Sundquist J, Memon AA. Thrombomodulin gene c.1418CT polymorphism and risk of recurrent venous thromboembolism. J Thromb Thrombolysis. 2016;42:135–141. doi: 10.1007/s11239-015-1328-x.

45. Manderstedt E, Halldén C, Lind-Halldén C, Elf J, Svensson PJ, Engström G, Melander O, Baras A, Lotta LA, Zöller B. Thrombomodulin (THBD) gene variants and thrombotic risk in a population-based cohort study. J Thromb Haemost. 2022;20:929–935. doi: 10.1111/jth.15630.

46. van der Velden PA, Krommenhoek-Van Es T, Allaart CF, Bertina RM, Reitsma PH. A frequent thrombomodulin amino acid dimorphism is not associated with thrombophilia. Thromb Haemost. 1991;65:511–513.

47. Aleksic N, Folsom AR, Cushman M, Heckbert SR, Tsai MY, Wu KK. Prospective study of the A455V polymorphism in the thrombomodulin gene, plasma thrombomodulin, and incidence of venous thromboembolism: the LITE Study. J Thromb Haemost. 2003;1:88–94. doi: 10.1046/j.1538-7836.2003.00029.x.

