## Supplemental Materials for "The Endothelium Modulates the Prothrombotic Phenotype of Factor V Leiden: Evidence from an Ex Vivo Model"

### Increased Activated Protein C Response to Thrombin Formation in Asymptomatic Factor V Leiden Carriers is Driven by the Endothelium: Evidence from and Endothelial Cell-based Ex Vivo Model

#### SUPPLEMENTAL METHODS

##### Materials

Human α-thrombin, protein C (PC) and activated PC (APC) were obtained from CellSystems (Troisdorf, Germany). Argatroban was obtained from Mitsubishi Pharma (Düsseldorf, Germany). Aprotinin and Streptavidin were purchased from PanReac AppliChem ITW Reagents (Darmstadt, Germany). Bivalirudin was obtained from The Medicines Company (Oxfordshire, UK). Biotinylated aptamers were synthesized and PAGE-purified by Microsynth (Balgach, Switzerland). Liquid Plate Sealer® and HRP Protector™ were purchased from Candor Bioscience (Wangen, Germany). The fluorogenic peptide substrate Boc-Asp(OBzl)-Pro-Arg-AMC (I-1560) was obtained from Bachem (Weil am Rhein, Germany) while the fluorogenic substrate Pyroglu-Pro-Arg-AMC (Pefafluor PCa) was purchased from Pentapharm (Basel, Switzerland). Batroxobin reagent was obtained from Siemens Healthineers (Marburg, Germany). Normal platelet-poor plasma was prepared in-house by pooling citrated plasma of at least four healthy blood donors. The tissue factor reagent for thrombin generation (PPP-reagent low) was purchased from Stago (Asnières-sur-Seine, France). Phospholipid emulsion containing phosphatidylserine, phosphatidylcholine and sphingomyelin was obtained from Rossix (Mölndal, Sweden). Ficoll Paque Plus was obtained from GE Healthcare (Solingen, Germany). Biotinylated bovine serum albumin (BSA), Dulbecco's phosphate-buffered saline (DPBS), ethylenediaminetetraacetic acid (EDTA), 0.5 % trypsin-EDTA solution (10x), glutaraldehyde 10 % solution, mouse IgG isotype control, fetal bovine serum (FBS), and Pierce™ BCA Protein Assay Kit were purchased from Thermo Fisher Scientific (Darmstadt, Germany). Dimethyl sulfoxide was purchased from WAK Chemie Medical (Steinbach, Germany). Rat tail collagen type I was purchased from Corning (Wiesbaden, Germany). EBM™-2 Basal Medium and EGM™-2 Endothelial Cell Growth Medium-2 BulletKit™ were obtained from Lonza (Basel, Switzerland). Russel's viper venom factor V activator was purchased from Loxo (Dossenheim, Germany). Mouse monoclonal antibodies against endothelial protein C receptor and thrombomodulin, and Janus Green cell normalization stain were purchased from abcam (Cambridge, United Kingdom). BSA and 3,3',5,5'-tetramethylbenzidine were obtained from Sigma Aldrich (St. Louis, United States). Fluorescently labelled, recombinant anti-human (REAfinity™) antibodies against CD31, CD309, CD201, CD141, CD34, and CD45 were obtained from Miltenyi Biotech (Bergisch Gladbach, Germany).

**Collection and Processing of Blood Samples**

Blood samples were obtained by venipuncture of an antecubital vein using 21-gauge winged infusion sets (Sarstedt, Nümbrecht, Germany). After discarding the first 2 mL, blood was drawn into EDTA tubes, lithium-heparin tubes (16 IU/mL), and citrate tubes (10.5 mmol/L, Sarstedt). Platelet-poor plasma was obtained from citrate tubes by centrifugation (2,600 x g, 10 minutes) within 30 minutes and stored at below -70°C until further processing.

**Genetic analysis**

Genomic DNA was isolated from EDTA blood using the Blood Core Kit (Qiagen, Hilden, Germany). The sequencing analyses were carried out on a Mini-Seq genome sequencer (Illumina, Santa Clara, CA, USA), which was used for next generation sequencing using a Nextera Rapid Capture Custom Enrichment (Illumina) including the following genes: *F2, F5, F7, F8, F9, F10, F11, F13A1, F13B, FGA* (fibrinogen alpha chain), *FGB* (fibrinogen beta chain), *FGG* (fibrinogen gamma chain), *VWF, GGCX, VKORC1, LMANT, MCFD2, SERPINC1* (antithrombin), *PROS1* (protein S), *PROC* (PC), *THBD* (thrombomodulin), and *PROCR* (endothelial protein C receptor, EPCR). Data were evaluated by SeqPilot (JSI medical systems, Ettenheim, Germany) software. For the description of sequence variations, the guidelines of the Human Genome Variation Society were applied, and variants were filtered according to minor allele frequency (MAF<1% in gnomAD).

**Quality Control of Cultured Endothelial Colony Forming Cells**

Cobblestone morphology of endothelial colony forming cells (ECFCs) was examined by light microscopy (Axiovert 25 or Axio Observer, both Carl Zeiss Microscopy, Oberkochen, Germany). Measurement of the total protein was performed amount using the Pierce™ BCA Protein Assay Kit (Thermo Fisher Scientific, Darmstadt, Germany) and staining of CD31 (platelet endothelial cell adhesion molecule, PECAM-1), CD309 (vascular endothelial growth factor receptor 2, VEGFR-2), CD201 (EPCR), and CD141 (thrombomodulin), CD34, and CD45 analyzed by flow cytometry as described elsewhere.[23] Briefly, cells were dissociated, resuspended in staining buffer (DPBS, pH 7, 2 % FBS, 0.5 mol/L EDTA), and $10^5$ cells were stained with fluorescently labelled antibodies for 30 minutes at room temperature in the dark. Cytometric measurements were performed using a Navios EX flow cytometer (Beckman Coulter Life Sciences, Brea, CA, USA) and analysis was performed using the FlowJo™ Software version 10.8 (BD Life Sciences, Ashland, USA).

**Aptamer-based Measurement of Thrombin and Activated Protein C**

Maxisorp Fluoronunc microtiter modules (Nunc A/S, Roskilde, Denmark) were coated with 10 µg/mL BSA-biotin, loaded with 10 µg/mL streptavidin, and blocked using 2 mg/ml BSA: Primed plates were treated with Liquid Plate Sealer® and stored in aluminium bags at 4°C until being used. For running the oligonucleotide-based enzyme capture assays (OECAs), pre-coated plates were incubated with 3'-biotinylated aptamers (HD1-22 for the thrombin-OECA or HS02-52G for the APC-OECA) and washed. Thrombin samples were further diluted 1:10 and diluted thrombin and APC samples were added to the aptamer-coated plates, respectively. After incubation and washing, detection of captured thrombin or APC was performed using the respective enzyme-specific fluorogenic peptide substrates (Boc-Asp(OBzl)-Pro-Arg-AMC for the thrombin-OECA or Pyroglu-Pro-Arg-AMC for the APC-OECA). Changes in fluorescence over time were measured using a fluorescence plate reader (Synergy 2, BioTek Instruments, Bad Friedrichshall, Germany). Calibration curves (in the same matrix as the samples) were processed in parallel covering a ½-log10 concentration range (0 to 10 ng/mL of thrombin, 0-272 pmol/L; or 0 to 50 ng/mL of APC, 0-910 pmol/L). Data obtained from the calibrators were interpolated by 4-parameter curve fit and used to calculate the thrombin or APC concentration in the samples.

**Assessment of Endothelial Cell-Dependent Activated Protein C Formation in a Purified System**

For generation of APC on ECFCs in a purified system, a buffer solution (10 mmol/L 4-(2-hydroxyethyl)-1-piperazineethanesulfonic acid, pH 7.4, 137 mmol/L NaCl, 4 mmol/L KCl, 11 mmol/L glucose, 2 mmol/L $CaCl_2$, 4 mg/mL BSA) was prepared. Cells were washed once with DPBS and once with buffer before addition of 50 nmol/L PC, and 0.1 U/mL thrombin in buffer (200 µL/well in 48-well plates). After one hour incubation at room temperature, the reaction was stopped, and generated APC was stabilized by diluting aliquots of the supernatant 1:10 into APC sample buffer. Samples were stored at below -70°C until APC was measured by OECA.

**Measurement of Coagulation Factors and Inhibitors in Plasma**

Plasma levels of factor II, factor V, factor VII, factor VIII, factor IX, factor X, factor XI, antithrombin, PC, and PS were measured on the Atellica® COAG 360 System using corresponding reagents (Siemens Healthineers, Erlangen, Germany).

#### SUPPLEMENTAL TABLES

**Table S1. Gene variants and range of activated protein C response in the autologous ex vivo model**

| | | **Non FVL *** | | **FVL VTE- †** | | **FVL VTE+ ‡** | |
|---|---|---|---|---|---|---|---|
| | | n | AUC APC/ AUC thrombin | n | AUC APC/ AUC thrombin | n | AUC APC/ AUC thrombin |
| ***F5*** **1691G>A (Factor V Leiden** | GG | 7 | 0.010-0.057 | 0 | - | 0 | - |
| | GA | 0 | - | 5 | 0.072-0.194 | 6 | 0.013-0.111 |
| | AA | 0 | - | 2 | 0.127-0.138 | 1 | 0.083 |
| ***F5*** **6755A>G (HR2 haplotype)** | AA | 7 | 0.010-0.057 | 6 | 0.072-0.194 | 5 | 0.016-0.111 |
| | AG | 0 | - | 1 | 0.153 | 2 | 0.028-0.100 |
| ***THBD*** **1418C>T** | CC | 4 | 0.047-0.057 | 3 | 0.072-0.194 | 5 | 0.016-0.111 |
| | CT | 3 | 0.010-0.031 | 1 | 0.093 | 1 | 0.013 |
| | TT | 0 | - | 3 | 0.127-0.153 | 1 | 0.027 |
| ***PROCR*** **4600A>G** | AA | 4 | 0.018-0.057 | 6 | 0.072-0.194 | 6 | 0.013-0.111 |
| | AG | 3 | 0.010-0.031 | 1 | 0.127 | 1 | 0.016 |
| ***PROCR*** **4678G>C** | GG | 2 | 0.047-0.057 | 3 | 0.093-0.153 | 3 | 0.013-0.111 |
| | CG | 4 | 0.010-0.055 | 3 | 0.072-0.194 | 2 | 0.027-0.100 |
| | CC | 1 | 0.031 | 1 | 0.149 | 2 | 0.028-0.083 |
| **All** | | 7 | 0.010-0.057 | 7 | 0.072-0.194 | 7 | 0.013-0.111 |

* In one subject (AUC APC/AUC thrombin = 0.031) the additional variants *THBD* 1477G>T and *F10* 424G>A were detected. † In one subject (AUC APC/AUC thrombin = 0.153) the additional variant *FGA* 1823G>C was detected. ‡ In one subject (AUC APC/AUC thrombin = 0.100) the additional variants *F13* 1730C>T and *FGB* 794C>T were detected. APC, activated protein C; AUC, area under the curve; FVL, factor V Leiden mutation; VTE, venous thromboembolism.

#### SUPPLEMENTAL FIGURES

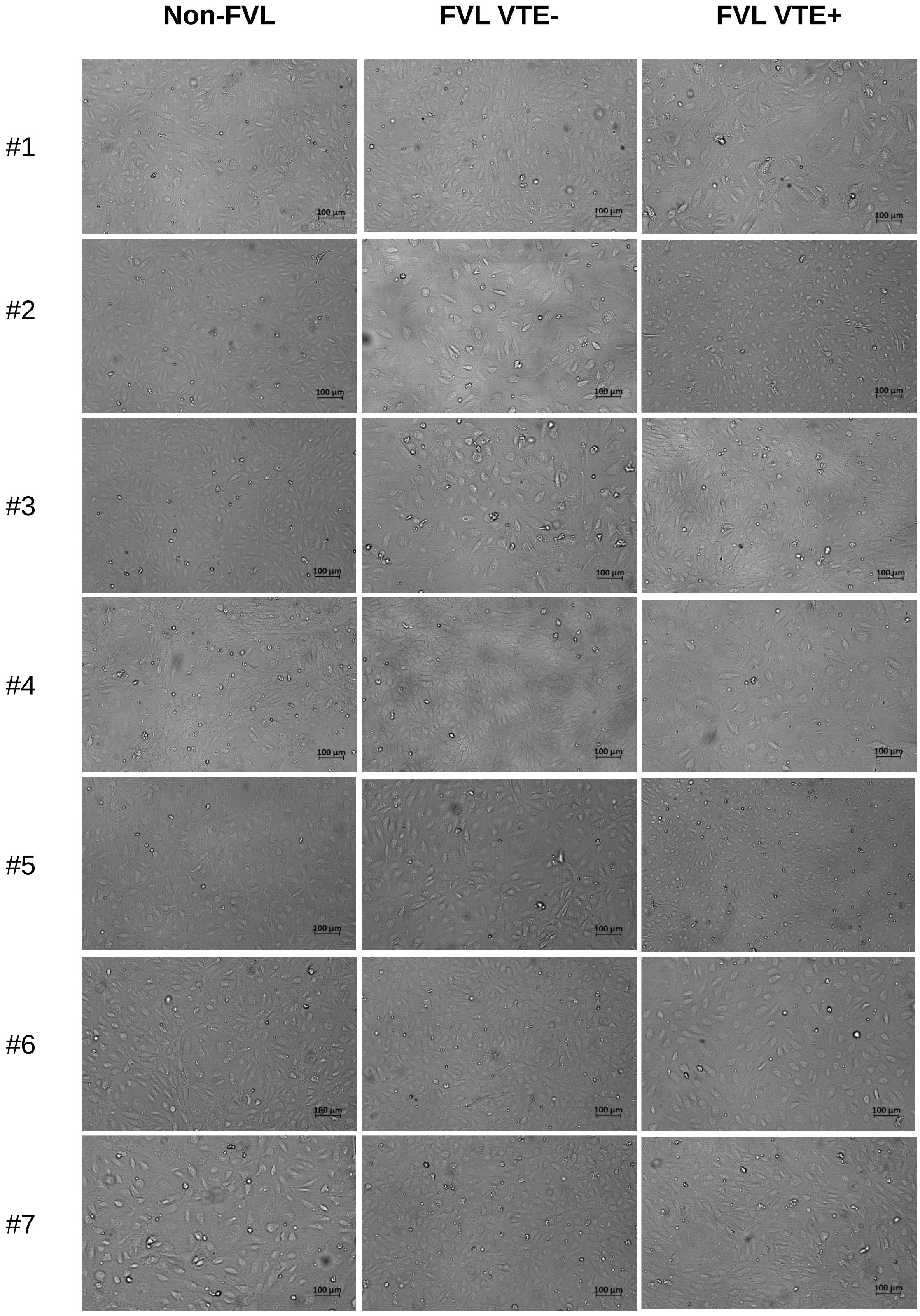


**Figure S1. Representative micrographs of endothelial colony forming cells.** Micrographs were obtained with an Axio Observer microscope using an Axiocam 702 mono camera (both Carl Zeiss Microscopy, Oberkochen, Germany). FVL, factor V Leiden; VTE, venous thromboembolism.

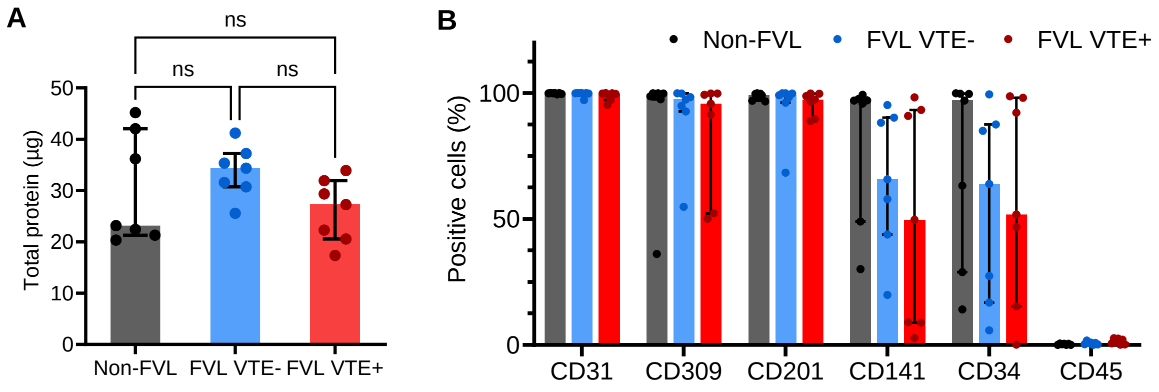


**Figure S2. Quality control of endothelial colony forming cells. A**, Cell lysates were obtained from confluent cell cultures on a 24-well plate using lysis buffer containing Triton X-100 (50 mM Tris HCl, 150 mM NaCl, 1 % Triton X-100, pH 8). Total protein amount was measured using the Pierce™ BCA Protein Assay Kit (Thermo Fisher Scientific, Darmstadt, Germany). **B**, Cells were dissociated, resuspended in staining buffer, and $10^5$ cells were stained with fluorescently labelled antibodies against CD31, CD309, CD201, CD141, CD34, and CD45 for 30 minutes. Cytometric measurements were performed using a Navios EX flow cytometer (Beckman Coulter Life Sciences, Brea, CA, USA) and analysis was performed using the FlowJoTM Software version 10.8 (BD Life Sciences, Ashland, USA). Data are shown as median and interquartile range. Differences between cohorts were assessed using the Kruskall-Wallis test followed by pairwise comparison using the Dunn procedure. The Bonferroni method was used to correct for multiple comparisons. No statistically significant differences between cohorts were observed. FVL, factor V Leiden mutation; ns, not significant; VTE venous thromboembolism.
